# Docosahexaenoic fatty acid-containing phospholipids affect plasma membrane susceptibility to disruption by bacterial toxin-induced macroapertures

**DOI:** 10.1101/2020.12.23.424114

**Authors:** Meng-Chen Tsai, Lucile Fleuriot, Sébastien Janel, David Gonzalez-Rodriguez, Camille Morel, Amel Mettouchi, Delphine Debayle, Stéphane Dallongeville, Jean-Christophe Olivo-Marin, Bruno Antonny, Frank Lafont, Emmanuel Lemichez, Hélène Barelli

## Abstract

Metabolic studies and animal knockout models point to the critical role of polyunsaturated docosahexaenoic acid (22:6, DHA)-containing phospholipids (PLs) in physiology. Here, we study the impact of DHA-PLs on the dynamics of transendothelial cell macroapertures (TEMs) tunnels triggered by the RhoA GTPase inhibitory exotoxin C3 from *Clostridium botulinum*. Through lipidomic analyses, we show that primary human umbilical vein endothelial cells (HUVECs) subjected to DHA-diet undergo a 6-fold DHA-PLs enrichment in plasma membrane at the expense of monounsaturated OA-PLs. In contrast, OA-diet had almost no effect on PLs composition. Consequently, DHA treatment increases the nucleation rate of TEMs by 2-fold that we ascribe to a reduction of cell thickness. We reveal that the global transcellular area of cells remains conserved through a reduction of the width and lifetime of TEMs. Altogether, we reveal a homeostasis between plasma membrane DHA-PLs content and large-scale membrane dynamics.

## Introduction

The plasma membrane attached to the cortical cytoskeleton forms a composite material that undergoes constant reshaping to perform essential cellular processes, including cell division, migration, phagocytosis and epithelial or endothelial semipermeable barrier organization and function (Levayer and Lecuit, 2012; Salbreux et al., 2012). Lipidomic approaches offer ways to quantitatively decipher the impact of fine-tuned changes in the composition of lipid acyl chains on membrane dynamics.

Phospholipids (PLs) often contain an unsaturated acyl chain at the sn-2 position that determines the biophysical properties of cellular membranes. Fatty acids (FA) are classified as saturated (S), monounsaturated (MU), and polyunsaturated (PU) by the number of double bonds present in the hydrocarbon acyl chain. Several glycerophospholipid classes, including phosphatidylcholine (PC), phosphatidylethanolamine (PE) and phosphatidylserine (PS), are the dominant constituents of the plasma membrane in addition to cholesterol (van Meer et al., 2008). Notably, phosphatidylcholine (PC) accounts for 40-50% of total phospholipids at the plasma membrane (van Meer et al., 2008). Variations in the length and number of double bonds in acyl chains lead to a remarkably large repertoire of phospholipid variants, such as PC(16:0/18:1, PE(18:0/20:4), and PS(18:0/22:6), conferring different biophysical properties, i.e., fluidity, packing order and curvature (Barelli and Antonny, 2016; Harayama and Riezman, 2018). The double bonds in polyunsaturated phospholipids allow acyl chains to twist at various angles, thereby providing the membrane with remarkably flexible properties (Manni et al., 2018). It is important to decode how the pattern of acyl chain variants in PLs translates into variations in cellular membrane dynamics (Harayama and Riezman, 2018; Pinot et al., 2014).

With 22 carbons and six double bonds, docosahexaenoic acid (DHA) is the most unsaturated form of the omega-3 fatty acids. Given the limited synthesis of this FA from linolenic acid, a dietary supply of DHA is essential to the functions of the retina and for spermatogenesis (Iizuka-Hishikawa et al., 2017; Shindou et al., 2017) in addition to brain function (Bazinet and Layé, 2014). In particular, animals fed with PUFA-free diets develop reduced visual functions paralleling the low DHA content in their retinas, outcomes that indicate the critical requirement of attaining DHA from the diet for visual function (Jeffrey and Neuringer, 2009). Lysophosphatidic acid acyltransferase 3 (LPAAT3), which catalyzes the esterification of DHA to generate lysophosphatidic acid and form precursors of PL, notably DHA-containing PC and PE, is particularly rich in the retina and testis (Yuki et al., 2009). Mice with LPAAT3 knocked out display male infertility and show visual impairment due to structural defects in the membranes of photoreceptors. Much remains to be learned on how DHA impacts the architecture and dynamics of the plasma membrane.

Recent works have shown that polyunsaturated lipids facilitate membrane processes requiring deformations at the nanometer scale. First, incorporation of polyunsaturated acyl chains into PLs facilitates endocytosis in model cellular systems and makes the pure lipid bilayer more flexible and prone to fission mediated by dynamin and endophilin (Pinot et al., 2014, Manni et al., 2018). These effects might explain why polyunsaturated phospholipids are necessary for proper synaptic vesicle formation (Tixier-Vidal et al., 1986). Second, polyunsaturated phosphatidic acid facilitates secretory granule exocytosis in neuroendocrine chromaffin cells, probably by stabilizing intermediates that contribute to a high-curvature membrane during fusion pore formation (Tanguy et al., 2020). Finally, polyunsaturated PLs modulate the activity of several mechanosensitive ion channels, including TRP, TRP-like and Piezo channels (Caires et al., 2017; Randall et al., 2015; Romero et al., 2019). Many of these effects have been proposed to arise from a reduction in the energetic cost of membrane bending and/or from a modulation of the energy required for protein conformational changes within the membrane matrix. However, whether and how the enrichment of cellular membranes with PUPLs modulates large-scale membrane dynamics remain to be elucidated.

Transcellular pores are observed in endothelial cell-lined vessels and form during the transcellular diapedesis of leukocytes (Aird, 2007; Braakman et al., 2016; Schimmel et al., 2017). Several toxins from pathogenic bacteria, such as RhoA-inhibitory exoenzymes from *Staphylococcus aureus* and *Clostridium botulinum*, can induce transendothelial cell macroaperture (TEM) tunnels (Lemichez et al., 2013). This TEM formation has been linked to increased vascular permeability and dissemination of *S. aureus* in tissues via the hematogenous route (Boyer et al., 2006; Munro et al., 2010; Rolando et al., 2009). Several bacteria secrete toxins that lower cell actomyosin contractility, thereby promoting cell spreading, which favors close contact between the apical and basal membranes and initiates their self-fusion (Boyer et al., 2006; Ng et al., 2017). The cellular dewetting physical model is based on the premise that spreading cells generate enough membrane tension for TEM nucleation and growth (Gonzalez-Rodriguez et al., 2012). Widening of TEMs is resisted by line tension, which is partially explained by the membrane curvature generated by torus-like pores (Gonzalez-Rodriguez et al., 2012; Stefani et al., 2017). After nucleation, an imbalance between the membrane and line tension causes TEMs to passively expand up to the maximal equilibrium size, which is stabilized by a newly formed stiff actomyosin cable that encircles TEMs (Stefani et al., 2017). TEMs are eventually sealed by active cytoskeleton-based processes (e.g., lamellipodia formation or purse-string contraction) (Maddugoda et al., 2011; Stefani et al., 2017). While considerable progress has been made in understanding the interactions between the membrane and actin cytoskeleton regulatory machinery in the control of TEMs, much remains to be known about the contribution of plasma membrane mechanical properties.

We investigated these areas by analyzing TEM dynamics in primary human endothelial cells subjected to polyunsaturated versus monounsaturated fatty acid diets. We show that membrane enrichment in DHA-containing phospholipids increases the probability of TEM nucleation while decreasing the width and the lifetime of transcellular tunnels.

## RESULTS

### Comprehensive analysis of the phospholipids in HUVECs fed with fatty acid diets

HUVECs represent a convenient endothelial cell model to study several aspects of TEM formation. We first determined whether these cells are amenable to defined fatty acid diets aimed at changing the acyl chain profile of their membrane phospholipids. We compared docosahexaenoic acid (DHA, C22:6) with oleic acid (OA, C18:1) diets, i.e., the most polyunsaturated acyl chain *versus* the most abundant monounsaturated acyl chain in PLs, respectively (Harayama and Riezman, 2018). HUVECs were first subjected to medium with lipoprotein-depleted serum (LPDS) followed by a diet of LPDS supplemented with bovine serum albumin (BSA) complexed either with docosahexaenoic acid (DHA, C22:6) or oleic acid (OA, C18:1). A comprehensive analysis of the lipidome of HUVECs subjected to the different diet conditions was conducted by comparing relative quantities of lipid species using a Q Exactive mass spectrometer.

While lipid starvation conditions decreased the triglyceride (TG) storing form of acyl chains, we did not detect significant changes in the relative distribution of phospholipid classes (Figure 1A and Sup. Figure 1A). We monitored the cellular lipidome of HUVECs fed with FA diets for different times. We recorded a massive increase in TG that peaked at 1 hour (Sup. Figure 2A). The incorporation of fatty acids into phospholipids occurred with slower kinetics, reaching a plateau at 6 hours, most notably for DHA incorporation into PC (Sup. Figure 2B). Importantly, the relative distribution of phospholipid classes was conserved between conditions, except for PE, which was reduced by 25% in the DHA-treated cells (Figure 1A).

**FIGURE 1.**
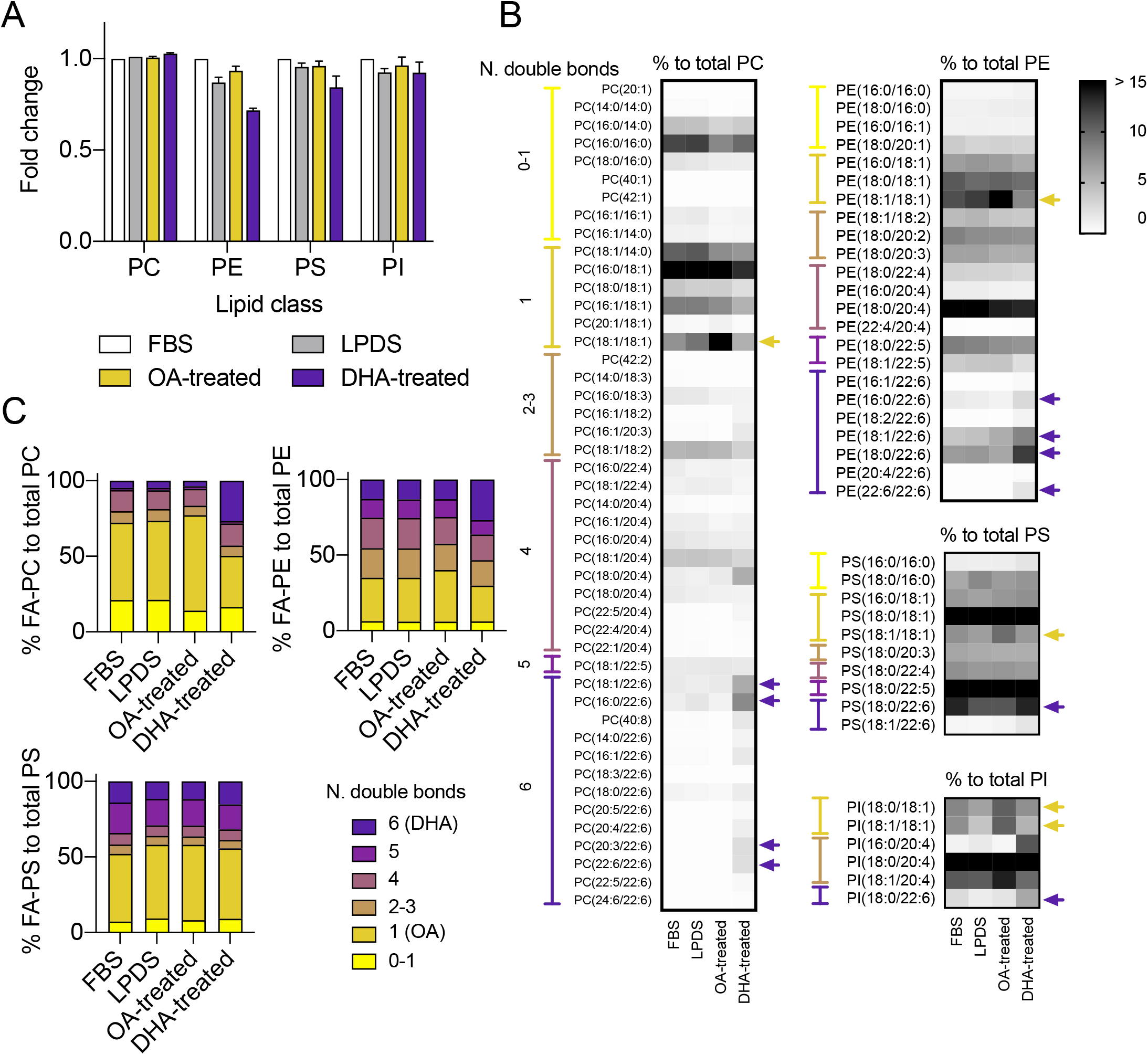
Analysis of phospholipid classes in HUVEC submitted to oleic acid (OA) *versus* docosahexaenoic acid (DHA). (A) Global fold change of phosphatidyl choline (PC), ethanolamine (PE), serine (PS) and inositol (PI) classes from HUVEC submitted to OA- or DHA-diet for 6 hours compared to FBS- or LPDS-cultured HUVEC. (B) Lipidic profiles comparison of the different PL classes: PC, PE, PS and PI upon OA-diet, DHA-diet compared to controls (FBS and LPDS). Values were normalized individually to the sum of each PL classes. (C) PL species distribution regarding their number of double bond (level of unsaturation): from zero to 6 double bonds as illustrated in figure 1B. AA and DHA contain 4 and 6 double bonds, respectively. LPDS = lipoprotein-depleted serum. (A-C) Data show means ± SEM; n>3 / experiments; 3 biological replicates.

As shown in Figure 1B and Sup1B, in contrast to OA, the DHA diet had an impact on the profile of phospholipids, which show enrichment in DHA-containing PL species (Figure 1B and Sup. Figure 1B). Specifically, DHA was incorporated in large amounts in PC and PE, with a 4-fold increase in PC(16:0/22:6) and a 2-fold increase in PE(18:0/22:6) (Figure 1B). In comparison, the remodeling of the anionic lipids PS and PI was modest, although we recorded an increase in PI(18:0/22:6) at the expense of PI(18:1/20:4), one of the major PI species. In sharp contrast, OA treatment had a narrow and slight impact on the acyl chain profile of phospholipids, inducing a specific increase in PL(18:1/18:1) at the cellular level, which was largely restricted to PC (Figure 1B). Overall, cells fed with DHA displayed considerable enrichment with polyunsaturated phospholipids, which we estimated as a polyunsaturated PC increase from 25% to 50%, at the expense of OA-containing PC (Figure 1C and Sup. Figure 2D). Furthermore, the addition of OA to HUVECs had a minor impact on the lipidome, which was already rich in OA-containing PLs and poor in DHA-containing lipids.

### Shifting the plasma membrane PL balance from the mono-unsaturated to the hexa-unsaturated form

The acyl chain profile of phospholipids varies according to subcellular localization (Antonny et al., 2015). Thus, we determined the impact of both FA diet conditions on the composition of PLs at the plasma membrane. For this experiment, we prepared giant plasma membrane vesicles (GPMVs) corresponding to plasma membrane blebs (Figure 2A). We observed the expected enrichment of the plasma membrane markers Annexin-V and Na^+^/K^+^ ATPase in the GPMV fractions (Figure 2B). Markers of internal compartments were observed in the total cell membrane fractions but were largely excluded from the GPMV fractions. The lipidomic analysis of the GPMVs compared to that of the total membrane fractions showed an enrichment in PS and sphingomyelin (SM), which are known to concentrate in the plasma membrane. In contrast, lipids characterizing membranes of internal compartments, such as diglyceride (DG; ER and lipid droplets) and lysobisphosphatidic acid (LBPA; late endosomes), were largely excluded from the GPMV fractions (Figure 2C). Moreover, quantitative analysis of PL classes in the GPMV fraction established the conservation of PL class distribution at the plasma membrane, including PE (Figure 2D).

**FIGURE 2.**
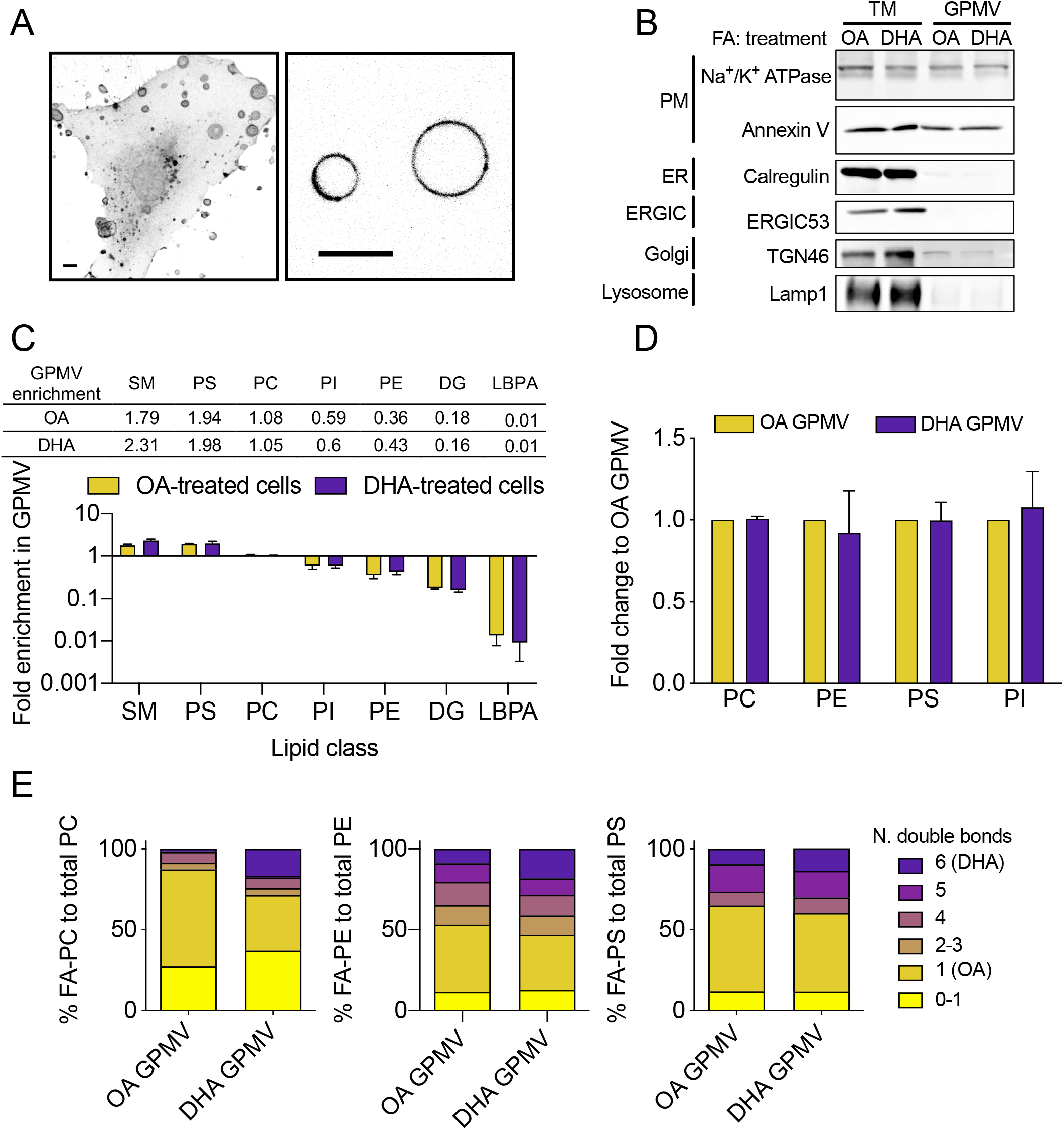
Lipidomic analysis of phospholipids from Giant Plasma Membrane Vesicle (GPMV). (A) 3D-projection of WGA-Alexa488 labeled GPMVs (left) and FM4-64 labelled GPMVs (right) from HUVEC. Scale bars 10 μm. (B) Western blot of total membrane (TM) and GPMV using different organelles markers. Plasma membrane (PM) marker: Na^+^/K^+^ATPase and Annexin V, Endoplasmic reticulum (ER) marker: Calregulin, ER-Golgi intermediate compartment (ERGIC) marker: ERGIC53, Golgi complex marker: Trans-Golgi Network 46 (TGN46), Lysosome marker: Lysosomal-associated membrane protein 1 (LAMP1). (C) Lipidomic analyses of GPMVs from OA- or DHA-treated cells. Lipid classes enrichment in GPMVs was calculated by dividing the relative content of each lipid class in GPMVs by those of total membranes. Value bigger than 1 indicate that lipids were enriched in GPMVs, value smaller than 1 indicate that lipids were excluded from GPMVs. SM, sphingomyelin; PC, phosphatidylcholine; PE, phosphatidylethanolamine; PS, phosphatidylserine; PI, phosphatidylinositol; DG, diacylglycerol; LBPA, lysobisphosphatidic acid. (D) Fold change of different PL classes in DHA-treated cells compared with OA-treated cells. (E) PL species distribution regarding their number of double bond (level of unsaturation): from zero to 6 double bonds as illustrated in figure 1B. AA and DHA contain 4 and 6 double bonds, respectively. (C-E) Data show means ± SEM; n>3 / experiments; 3 biological replicates.

We analyzed the changes in the acyl chain composition of PLs in the plasma membrane-derived GPMVs prepared from cells subjected to the two fatty acid diets. As observed for the total membrane fraction, the GPMV fraction from DHA-treated cells was enriched in DHA-containing PC, the dominant PL class, by 10-fold (from 1.6% to 16.7%) and PE species by 2-fold (from 8.8% to 18.2%) at the expense of monounsaturated species as compared to GPMV from OA-treated cells (Figure 2E). Thus, the DHA diet triggered a 1.9-fold reduction in OA-containing PC compared to OA-treated cells. Interestingly, these variations are accompanied by an increase in saturated PC species in agreement with recent work by Levental et. al. (Levental et. al., 2020).

Our comprehensive analysis of the HUVEC lipidome establishes that these cells have a plasma membrane intrinsically rich in OA-containing PLs, a profile that can be largely shifted to polyunsaturated DHA-containing PLs upon exposure to a high-DHA fatty acid diet.

### The DHA diet leads to smaller pores in the TEM population

Inhibition of the small GTPase RhoA by bacterial ExoC3-like toxins induces the nucleation and expansion of TEMs (Boyer et al., 2006). This effect is triggered by a collapse of RhoA-driven actomyosin contractility and leads to cell spreading and a reduction in cell thickness (Figure 3A)(Ng et al., 2017). We first verified the absence of the impact of RhoA signaling shutdown on the proper incorporation of OA or DHA into PLs. To this end, HUVECs were incubated in LPDS medium as the sole treatment or in LPDS containing the RhoA-inhibitory C3-exoenzyme (ExoC3). Next, the cells were incubated for 1 to 6 hours in LPDS medium supplemented with bovine serum albumin (BSA) complexed with either OA or DHA fatty acids. We observed that ExoC3 treatment did not interfere with the enrichment of DHA-PL or OA-PL species in cellular membranes (Sup. Figure 3), suggesting that RhoA signaling did not impact the metabolism of OA/DHA acyl chains in a manner that would interfere their proper incorporation into PLs species. In parallel, we measured similar level of RhoA ADP-ribosylation in cells subjected to OA and DHA fatty acid diets (data not shown) and verified the proper disruption of actin stress fibers due to ExoC3 action under the different diet conditions (Figure 3B).

**FIGURE 3.**
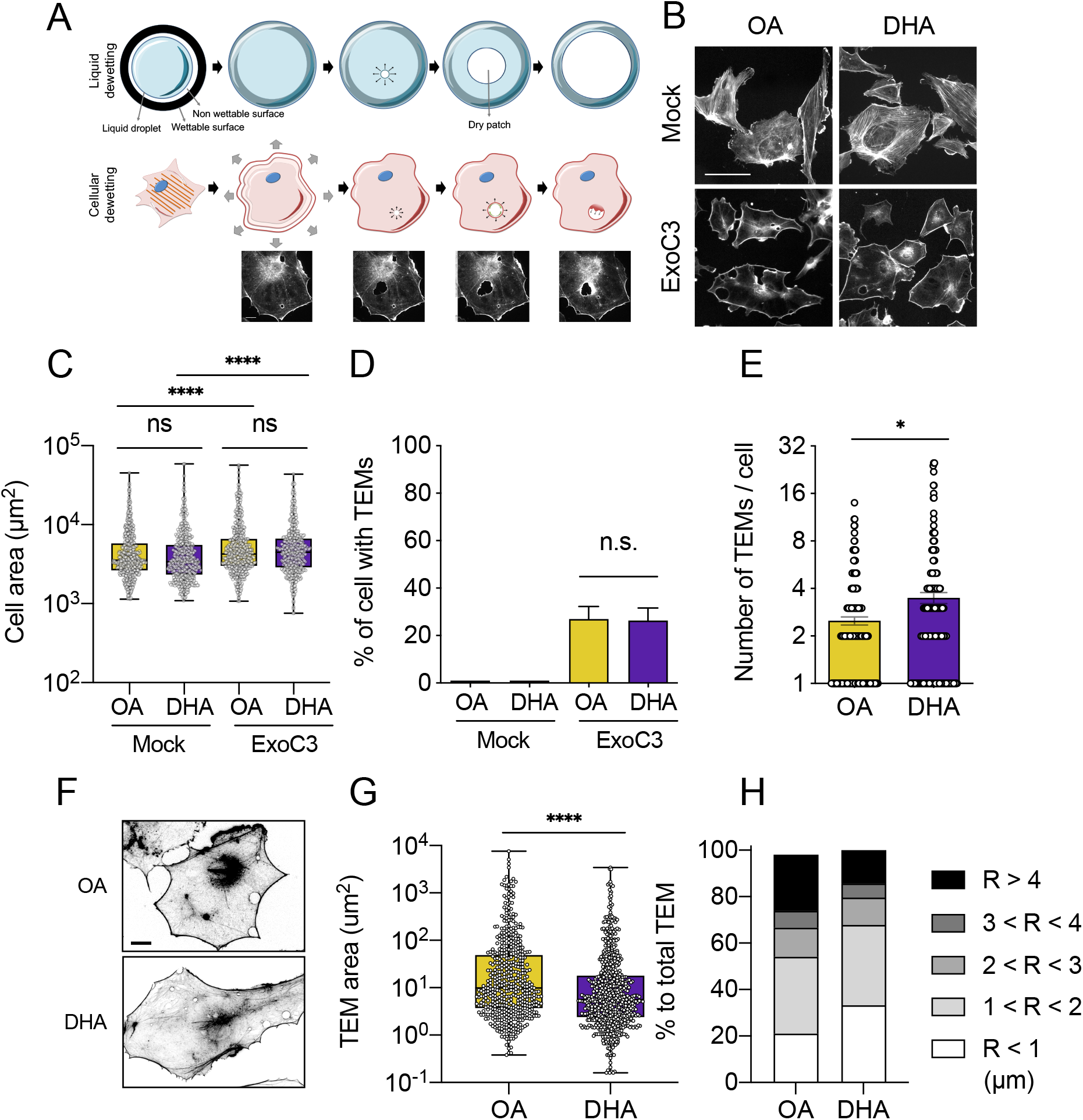
The impact of DHA on TEM parameters. (A) Schematic representations of liquid dewetting physics phenomenon and TransEndothelial cell Macroaperture dynamics. Scale bar 20 µm. (B-H) HUVEC were treated with C3 exoenzyme or without (Mock) for 16 hours prior to 6-hours fatty acid diet (OA or DHA) as sole source of exogenously added acyl chain. (B) FITC-phalloidin staining of HUVEC in Mock, OA or DHA conditions. Scale bar 100 µm. (C) Cell area of cells on OA- or DHA-diet. Data show median ± max to min; cells > 450 / experiments; 2 biological replicates. (D) Percentage of HUVEC with at least one TEM in the population in OA or DHA conditions. Data show means ± SEM; cells > 200 / experiments; 3 biological replicates. (E) Number of TEMs per cell under OA- or DHA-diet. Data show median ± max to min of >150 cells from 3 independent experiments (> 50 cells / experiment). (F) Representative FITC-phalloidin staining of OA or DHA-treated cells intoxicated with C3 toxins. Scale bar 20 µm. (G) Graph shows median values of TEM area in fixed cells treated with either OA or DHA. Data show median ± max to min of > 450 TEMs from 3 independent experiments (>40 cells/ experiments). (H) Distribution of TEM sizes in HUVEC under OA- or DHA-diet. (C, D, E, G) Data were analyzed with nonparametric Mann-Whitney statistical test. Data are significant with p < 0.05(*), p < 0.0001 (****) or not significant, ns.

We analyzed the impact of OA or DHA treatment on the spreading of cells induced by ExoC3. Measures of cell area showed no significant difference between the two diet conditions in non-intoxicated cells (Figure 3C). When cells were treated with ExoC3, we recorded a 1.2-fold spreading of both OA- and DHA-treated cells, indicating that cell enrichment in DHA-containing PL did not significantly influence the extent of HUVECs spreading in response to the inhibition of RhoA.

Next, we analyzed TEM parameters on fixed cells stained with FITC-phalloidin to label filamentous actin accumulating around TEMs. We observed that approximately 25% of cells displayed at least one TEM with no significant difference between the cells cultured under the two fatty acid diet conditions (Figure 3D). In this subpopulation, we recorded a net increase in the density of TEMs in the DHA-treated cells of 3.5 ± 0.3 TEM/cell *versus* 2.5 ± 0.1 TEM/cell for the OA-fed cells (Figure 3E). In addition, we observed a 1.6-fold decrease in the TEM area in the DHA-treated cells compared with the OA-treated cells, with A_-DHA_ = 5.6 μm^2^ *versus* A_-OA_ = 9.9 μm^2^, respectively (Figure 3F and 3G). Despite the moderate global effect of DHA on TEM median size, a thorough analysis of the distribution of TEMs revealed that the DHA diet induced a major shift toward TEMs of small size (R<1 µm; 21% to 33%) at the expense of large TEMs (R>4 µm; 24% to 14%) (Figure 3H). Altogether, these results indicated that inhibition of RhoA and incorporation of DHA into phospholipids did not interfere with each other, while DHA-PL enrichment in the plasma membrane led to an increase in TEM density and a decrease in TEM size.

### The DHA diet increases TEM nucleation frequency

TEM tunnels form labile openings (Figure 4A) (Video 1). After nucleation and growth, TEMs reach a stable state in which they oscillate around a maximal area. After this period of latency, TEMs undergo a phase of closure via actin-dependent processes involving either purse-string contraction or membrane wave extension (Maddugoda et al., 2011). Here, we noticed that approximately 70% of the TEMs resealed via a purse-string contraction phenomenon regardless of the fatty acid diet (not shown).

**FIGURE 4.**
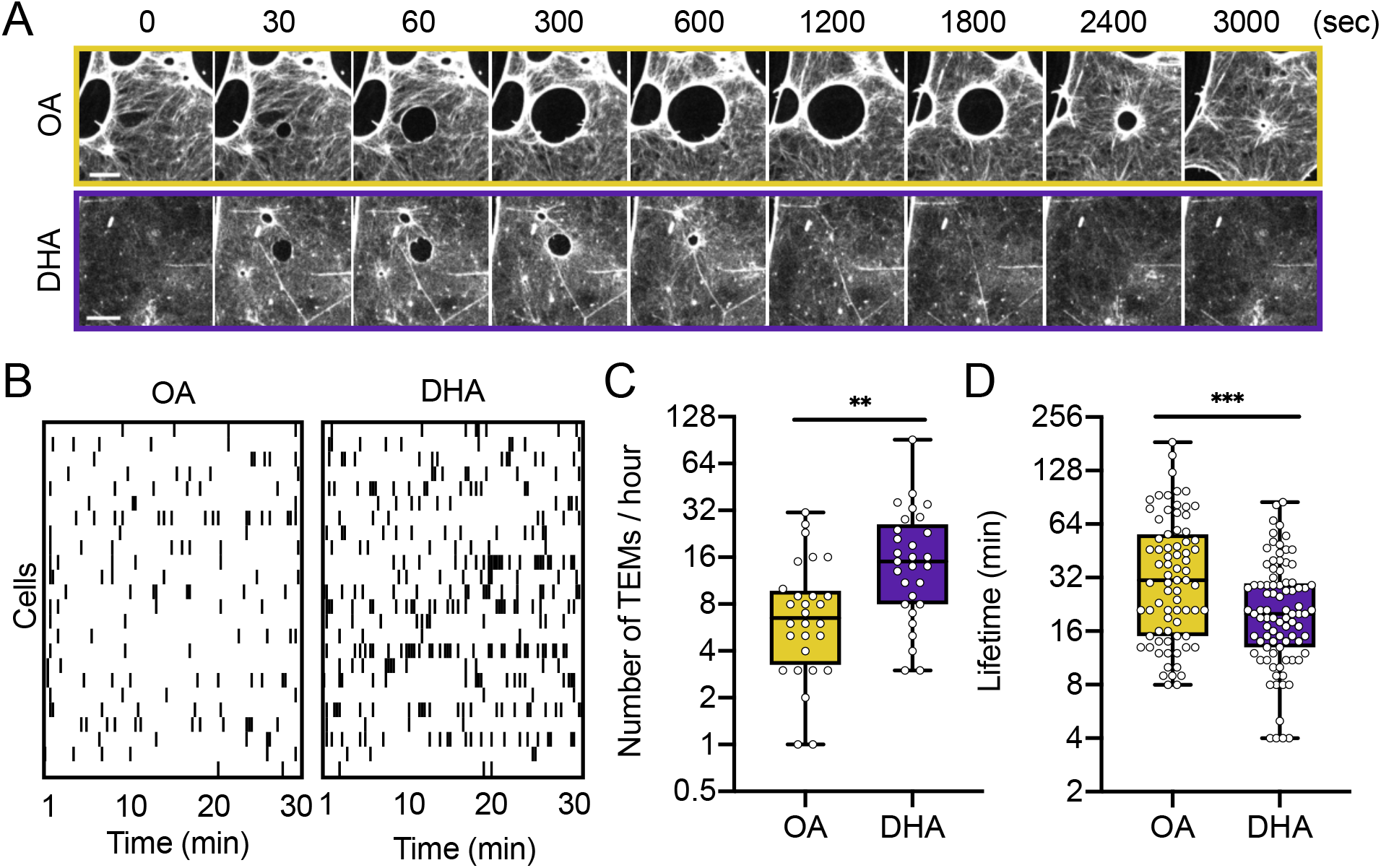
Impact of DHA-enriched membrane on TEM opening dynamics. Time-lapse images of TEM opening dynamics in OA- or DHA-fed cells using LifAct-GFP as label. Scale bars 10 µm. (B-C) Frequency of TEM opening events per cell in cells treated with ExoC3 and fed with either OA or DHA. (B) Each row on the Y-axis of the diagram pinpoints all opening events of TEM in a single cell, each black bar is an individual opening event. (C) Graph shows total number of TEM opening events per cell per hour. Data show are median ± max to min; n>28 cells from 28 independent experiments. (D) Graph shows the distribution of values of TEM opening lifetimes measured for each TEM. Data are median ± max to min, n>75 TEMs from >8 independent experiments. (C-D) Data were analyzed with nonparametric Mann-Whitney statistical test and are significant with p < 0.01(**) and p < 0.0001.

To quantitatively analyze TEM dynamics, we recorded the cycles of TEM opening and closing by time-lapse video in LifeAct-GFP-expressing cells, allowing us to determine the frequency of TEM nucleation and their complete lifetime. Interestingly, this analysis revealed a critical impact of the DHA fatty acid diet. Figure 4B shows that the DHA-fed cells had a higher frequency of opening events during the recording period than OA-fed cells (Video 2 and 3). Mean values were *N*=19.3 events/h for the DHA-fed cells *versus N*=8.9 events/h for the OA-fed cells (Figure 4C). Moreover, we measured that the lifetime for complete TEM opening and closing cycle was 1.7-fold shorter in the DHA-fed cells than in OA-fed cells, i.e., mean values of 24.6 ± 1.8 min for the DHA-fed cells *versus* 42.0 ± 4.0 min for the OA-fed cells (Figure 4D). TEM cycles encompass two dynamic phases of opening and closure and a phase of latency where the TEM area oscillates within approximately 5% of its maximal size. While the time for TEM opening was not affected by DHA, we recorded a decrease in the duration of both the latency and closure phases (Table 1). The probability of observing a TEM in a cell depends on the product of TEM nucleation frequency by their lifetime. The net consequences of reducing the TEM lifetime and increasing the nucleation rate in DHA-treated cells explain the minimal difference in TEM density between the two conditions. We concluded that membrane enrichment in DHA-PLs increases the nucleation rate of TEMs with short lifetimes.

**Table 1.** Effect of FA diet on TEM parameters

|  | OA | DHA | Statistics |
| --- | --- | --- | --- |
| Cell area ($\mu\text{m}^2$) | 6330 ± 271 | 6003 ± 251 | ns |
| Cell thickness (nm) | 190 ± 6 | 164 ± 6 | ** |
| % of cells with TEMs | 27 ± 3 | 26 ± 3 | ns |
| Number of TEMs/cell | 2.5 ± 0.1 | 3.5 ± 0.3 | * |
| TEM frequency (TEMs/hour) | 8.9 ± 1.4 | 19.3 ± 3.2 | ** |
| TEM max size ($\mu\text{m}^2$)<sup>a</sup> | 37.4 (27.6-59.4) | 25.4 (21.2-28.6) | * |
| Opening time (sec) | 179 ± 21 | 162 ± 23 | ns |
| Initial opening speed ($\mu\text{m}^2/\text{sec}$)<sup>a</sup> | 0.47 (0.35-0.95) | 0.39 (0.29-0.54) | ns |
| Late opening speed ($\mu\text{m}^2/\text{sec}$)<sup>a</sup> | 0.38 (0.22-0.61) | 0.27 (0.14-0.37) | * |
| Latency phase (min) | 7.3 ± 0.6 | 5.3 ± 0.4 | * |
| Closure time (min) | 16.3 ± 1.4 | 11.1 ± 0.8 | ** |
| Closure speed ($\mu\text{m}^2/\text{sec}$)<sup>a</sup> | 4.3 (3.4-6.1) | 2.7 (2.1-3.3) | *** |
| Life time (min) | 42.0 ± 4.0 | 24.6 ± 1.8 | *** |

Data show the means ± SEM; data were analyzed by Mann-Whitney test
<sup>a</sup> data show the median (95% CI, upper-lower)
ns, not significant; \* $p$ <0.05, \*\* $p$ <0.01, \*\*\* $p$ <0.001

### The DHA diet decreases cell thickness

Using atomic force microscopy (AFM), we measured the elasticity of cells, which is mainly affected by cortical actin (Figure 5A). RhoA inhibition greatly reduced cell elasticity, by 1.7-fold, as reported (Ng et al., 2017). However, we measured similar elasticity in the cells subjected to the OA and DHA treatments (5.1 ± 0.2 kPa *versus* 5.8 ± 0.3 kPa, respectively) (Figure 5B-C), suggesting negligible impact of DHA enrichment on cortical elasticity.

**FIGURE 5.**
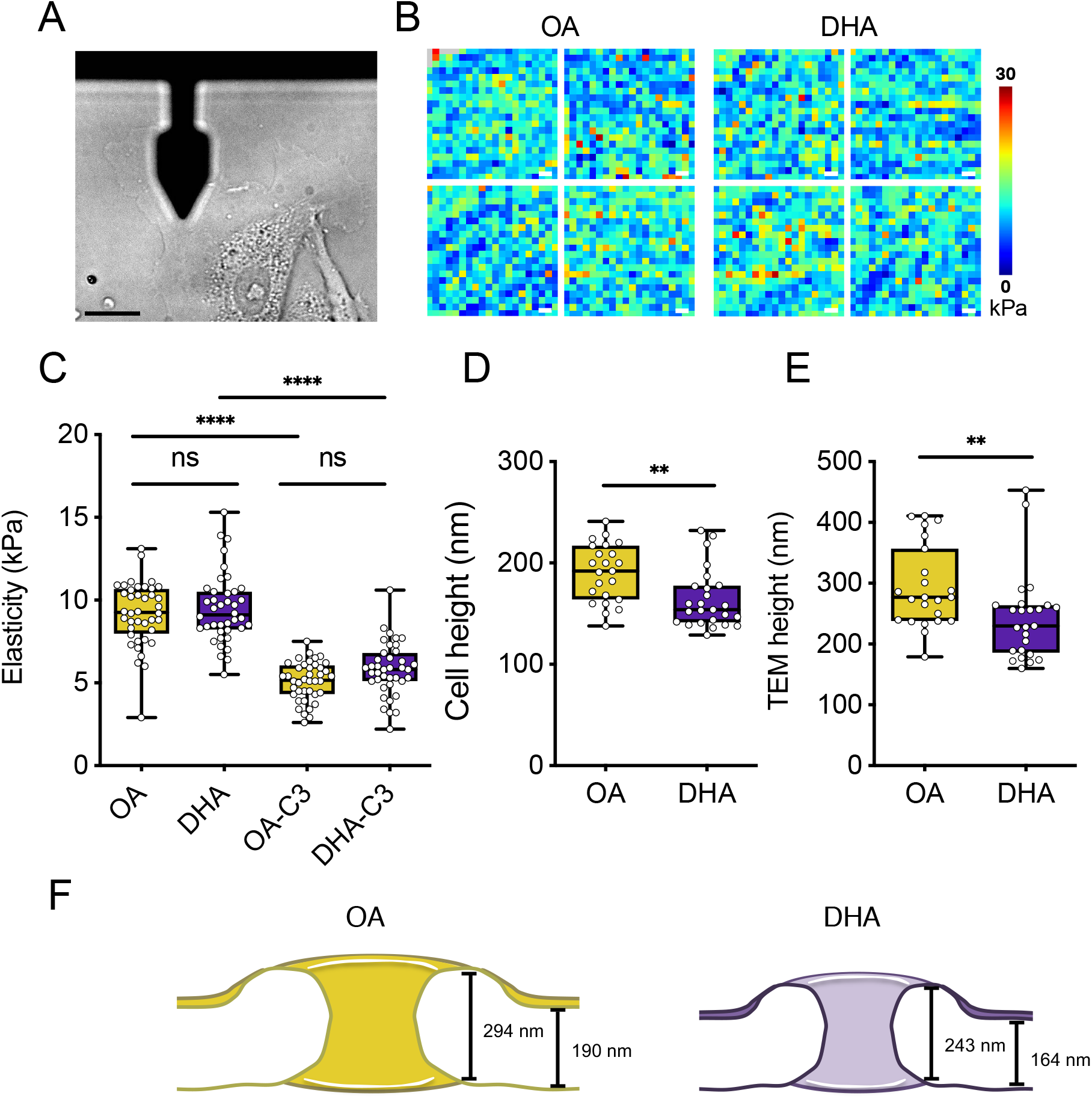
Impact of DHA on cell geometry. (A) Atomic force microscopy (cantilever shape seen in forground) is used to measure the mechanical parameters of the cell (background in bright field). Scale bar 20 µm. (B) Heatmap of cell elasticity on OA- and DHA-treated cells. Scale: 10 µm^2^ per field of view. (C) Graph shows the elasticity of Mock or C3-intoxicated cells treated with OA or DHA. Data show median ± max to min; n = 40 TEMs. Data were analyzed with one-way ANOVA with Bonferroni correction and are significant with ****p < 0.0001 or not significant, ns. Thickness of the cell at the periphery of TEM (D) and thickness of TEM border (E) were measured by AFM using zero-force topology. (F) Graph shows the curvature of TEMs derived from the TEM height measurements. (D-F) Data show median ± max to min; n > 20 TEMs. Data were analyzed with Mann-Whitney test and are significant with **p < 0.01.

On the other hand, we measured differences in the topology of the TEMs with AFM by scanning the cell surface with zero force. Interestingly, DHA treatment decreased the cell thickness to 1.2-fold; i.e., mean values for the DHA-fed cells was 164 ± 6 nm *versus* 190 ± 6 nm for the OA-fed cells (Figure 5D). As previously reported, the TEM rim is elevated and forms a ridge structure (Maddugoda et al., 2011). In accordance with the measures of cell thickness, the height of the TEM ridge was decreased 1.2-fold in the DHA-enriched cells (Figure 5E). We concluded that DHA enrichment decreases the thickness of TEMs without affecting the cell elasticity contributed by the cellular cortex (Figure 5F).

### The impact of DHA-PL enrichment on TEM opening kinetics

The maximal area reached by TEMs depends on two parameters: the relaxation of membrane tension as TEMs open, which controls their opening speed, and the time required for a cell to encircle TEMs with a stiff actomyosin cable that prevents further widening (Stefani et al., 2017). Dynamic parameters were analyzed with a custom-made Icy-based program that automatically segments the LifeAct-GFP-decorated actin-rich circumference of TEMs as a function of time (Figures 4A and 6A). This analysis enabled us to determine the maximal area and opening and closing speeds.

Interestingly, we observed that, under LPDS conditions, approximately 11% of the TEMs in the OA-treated cells *versus* 3% of the TEMs in the DHA-treated cells resumed their enlargement 104 ± 33 seconds after stabilization, i.e., after they had reached the first stable state (Sup. Figure 4), suggesting that DHA-PL-rich membranes form pores with greater stability. In parallel, we performed a comparative super-resolution stimulated emission depletion (STED) microscopy analysis of the actin structures around the TEMs in the OA- and DHA-fed cells; i.e., we examined the typical actomyosin belt and membrane wave-containing dendritic F-actin network (Stefani et al., 2017). No significant difference in actin organization around the TEMs between the two conditions was recorded (Figure 6B). For the quantitative analysis, we focused on the first equilibrium state reached by the TEMs. We recorded and defined the median size of the TEMs over the recording period (Figure 6C). We also determined the initial speed of TEM opening, which correlates with membrane tension that drives the opening of TEMs (Gonzalez-Rodriguez et al., 2012). A comparative analysis of the initial speeds of opening between 10 and 20 seconds in the OA- and DHA-fed cells revealed no significant difference between the DHA-fed cells (V_i-DHA_ = 0.39 μm^2^ s^-1^) and OA-fed cells (V_i-OA_ = 0.47 μm^2^ s^-1^) (Table 1). In contrast, when we compared the median opening speed over the first 20-70 seconds, the values were 1.4-fold lower for the DHA-fed cells (V_0-DHA_ = 0.27 μm^2^ s^-1^) than for the OA-fed cells (V_0-OA_ = 0.38 μm^2^ s^-1^) (Figure 6D). In parallel, we assessed the maximal size of the TEMs during the recording periods. Consistent with measures of the fixed cells, the TEM maximal size was increased by approximately 1.5-fold in the DHA-treated cells compared with the OA-treated cells, with S_max-DHA_ = 25 ± 2 μm^2^ *versus* S_max-OA_ = 37 ± 7 μm^2^ (Figure 6E). In accordance with the TEM size of the fixed cells, we recorded, in live cells, an increase in small (R<3 µm) TEMs from 38% to 60% at the expense of large (R>4 µm) TEMs (from 42% to 20%) in the DHA-treated cells *vs* OA-treated cells (Figure 6F). Consistent with the significant decrease in TEM size and in the speed of opening recorded for the DHA-fed cells, we observed no difference in the time to reach the maximal surface area, i.e., 179 ± 21 seconds for the OA-fed cells and 162 ± 23 seconds for the DHA-fed cells (Figure 6G). Taken together, our data establish that, despite having no effect on the initial speed of opening, DHA-PLs reduced the overall TEM opening speed, thereby impacting the maximal TEM size.

**FIGURE 6.**
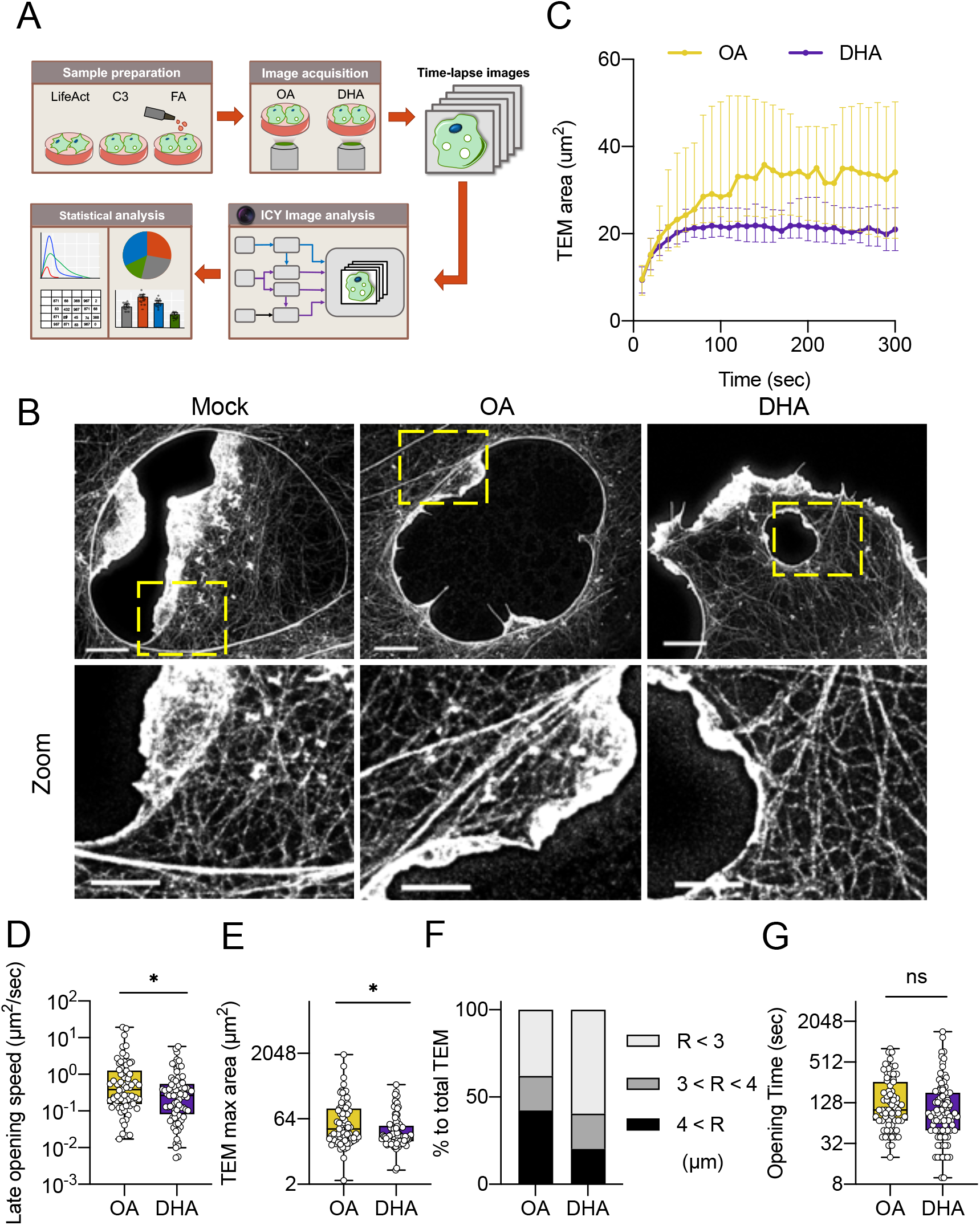
Impact of DHA-enrichment on TEM opening parameters. (A) Experimental workflow. HUVEC cultivated in LPDS medium were transfected with LifeAct-GFP, treated with C3 toxin, and fed with OA- or DHA-BSA for 6 hours. The dynamics of each TEM was recorded by spinning disk confocal microscopy and was analyzed through custom made Icy-based protocol. (B) Super-resolution stimulated emission depletion (STED) microscopy images of phalloidin-StarRed in C3 intoxicated cells treated without any fatty diet (Mock) or with OA- or DHA-diet. Scale bar 5 (upper) and 2 (lower) µm. (C) Graph shows median values of TEM area as a function of time in cells treated with either OA (blue, n=71 TEMs) or DHA (pink, n=94 TEMs) as sole source of exogenously added acyl chain. Values correspond to average surface of TEMs. Data show median ± 95% CI from 28 independent experiments. (D) Distribution of opening speed of TEM between 20-70 seconds. (E) Graph shows median values of TEM maximum area in cells treated with either OA or DHA. (F) Distribution of TEM sizes in HUVEC under OA- or DHA-diet. (G) Graph shows distribution of time durations to reach 95% of maximum area. (D, E, G) Data show median ± max to min, n>70 TEMs for each condition from 28 independent experiments. Data were analyzed with Mann-Whitney test and are significant with *p < 0.05 or not significant, ns.

The cellular dewetting physical model, which describes TEM dynamics (Gonzalez-Rodriguez et al., 2012; Stefani et al., 2017), indicates that the initial TEM opening speed (at the very first opening stage) is proportional to the membrane tension. Because the initial speed was similar between the OA- and DHA-treated cells, the average cell membrane tension is expected to be comparable between the two conditions. This conclusion was supported by the observation of comparable spreading of the OA- and DHA-fed cells because the cell spreading area correlates with cell membrane tension.

According to the physical model, an estimate of the global membrane bending rigidity is given by

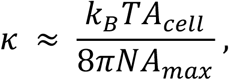

where *N* is the number of TEMs simultaneously evident per cell, *A*_max_ is the maximum area of a TEM, *k*_*B*_ is the Boltzmann constant, *T* is the temperature, and *A*_cell_ is the total cell spreading area. This equation is based on the relaxation of membrane tension during TEM opening, as described by Helfrich’s law (Helfrich, 1973). Membrane relaxation is larger when the bending rigidity κ is larger. In deriving this equation, we assume a joint effect of *N*, the number of simultaneous TEMs on membrane, on tension relaxation, which affects each TEM. Because the cell spreading area *A*_cell_ is the same in the OA- and DHA-fed conditions, this equation indicates that the global bending rigidity κ is inversely proportional to the product *N*·*A*_max_. A comparison between the two conditions leads to

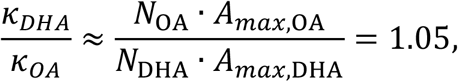

whose difference from 1 is not statistically significant. Thus, this calculation suggests that the average bending rigidity at the cellular scale is similar between cells in the OA- and DHA-fed conditions.

Overall, the interpretation of the experimentally observed TEM dynamics in light of the physical model suggests that average tension and bending rigidity of the membrane in association to cortex composite material are unchanged in cells subjected to these OA and DHA treatments. This result points to a regulation mechanism of global membrane dynamics controlling TEM opening that is robust despite changes in membrane lipid composition, which dictate the TEM nucleation rate.

## Discussion

Since the discovery of the cellular dewetting phenomenon, the contribution of plasma membrane mechanical properties to the dynamics of TEMs remains to be elucidated. Here, we show that the DHA fatty acid diet induces a shift in the acyl chain composition of phospholipids at the plasma membrane of endothelial cells, with an increase in DHA-PLs, and greatly affects TEM dynamics. Remarkably, DHA-PLs enrichment changes membrane dynamics, i.e., nucleation and lifetime of TEMs, in a coordinated manner to ensure the relative conservation of the overall TEM width per cell, and shifts the size range of the TEMs to a smaller range via reduction of opening speed. Moreover, DHA-PL enrichment reduced uncontrolled resume TEM growth. Collectively, these findings indicate that DHA-PL facilitates the nucleation of smaller TEMs displaying shorter lifetimes. Conversely, deficient DHA-PLs may lead to the opening of unstable and wider TEMs.

DHA-PL enrichment at the plasma membrane leads to a decrease in TEM maximal size and an increase in the number of simultaneous TEMs present in a cell. Strikingly, the total maximum TEM area, obtained as the product of the maximum area per TEM by the number of TEMs, remains constant between OA- and DHA-fed cells. Together with the physical interpretation provided by the cell dewetting model (Boyer et al., 2006; Gonzalez-Rodriguez et al., 2012), this observation suggests a conservation of the global membrane mechanical response at the scale of the entire cell. Therefore, whereas DHA-PL enrichment promotes the frequency of TEM nucleation events by 2-fold, the regulation of membrane mechanical characteristics at the scale of the whole cell appears sufficiently robust for the cell to cope with these changes, leading to a conserved total TEM area over the cell. To maintain the overall conservation of the TEM area, DHA-PL-enriched cells compensate for the increase in the nucleation rate by reducing the TEM opening speed and thereby maximal size. Moreover, we record a decrease of lifetime that is ascribed to a reduction in both the phase of latency and closure without affecting the initial opening phase.

Enrichment of DHA-PLs at the plasma membrane decreases the speed of opening, while the time to reach the maximal size is not affected. Consistent with the conserved time frame of TEM opening, we observed that actin organization around TEMs formed in the DHA- and OA-fed cells showed no significant difference. Nevertheless, we observed that TEMs were less stable in the OA-fed cells and were more prone to resume their enlargement. Lower TEM stability may reflect defects in the recruitment or activity of actin-crosslinking proteins around the edge of the TEMs that are yet to be identified. Importantly, we previously reported that the absence of TEM stabilization is linked to massive hemorrhage induced by an ExoC3-chimeric toxin derived from the *B. anthracis* lethal toxin (Rolando et al., 2009). This provides additional evidence that DHA might play a key function in the appropriate homeostasis of the endothelial barrier, pointing to a likely role in stabilizing large pore structures that are observed along the vascular system (see for review) (Aird, 2007; Lemichez et al., 2010).

Finally, we found a decrease in cell thickness in the DHA-PL-enriched cells. Applying force to the apical part of the plasma membrane of normal growing cells is sufficient to bring membranes in close apposition and trigger the nucleation and opening of transcellular pores (Ng et al., 2017). Consistent with this idea, HUVECs intoxicated with ExoC3 were thinner than the control cells, i.e., with medians near the edge of the cells at 332 nm *versus* 462 nm, respectively (Ng et al., 2017). It is therefore reasonable to think that enriching DHA-PLs may enhance close plasma membrane apposition and increase the probability of pore nucleation by reducing the energy cost to initiate a fusion event. As discussed below, changes in the asymmetric distribution of polyunsaturated phospholipids in DHA-PL-enriched cells might contribute to membrane apposition and fusion.

The plasma membrane is asymmetric in both lipid classes and lipid unsaturation (Lorent, et al., 2020). The outer leaflet contains mostly PC and SM, whereas the inner leaflet contains 3 major PL classes, namely, PC, PE and PS. Coarse-grained molecular dynamics simulations on asymmetric phospholipid bilayers show that DHA-PLs facilitate membrane tubulation only when they are located on the convex side of the deformation (Tiberti et al., 2020). This effect is due to the ability of DHA-PLs to switch between several twisted conformations in a convex environment, notably to adopt a conformation in which the polyunsaturated acyl chain occupies voids between polar heads and invades the water-lipid interface. DHA-fed cells showed a 10-fold higher content of DHA-PC species at the plasma membrane than was evident in the OA-fed cells (16.7 and 1.6%, respectively). In contrast, the amount of DHA-PE and DHA-PS species was only approximately 2-fold higher in the DHA-fed HUVECs (18.2% and 14.1%, respectively) than in OA-fed cells (8.8% and 9.5%, respectively). Consequently, the DHA diet might not only increase the overall DHA-PL content of the plasma membrane but might also reduce DHA asymmetry because PC is the most affected PL and is quite evenly distributed between the two leaflets. Such a change coupled with the lack of contractile forces mediated by the cytoskeleton, due to the ExoC3 effect, may favor large undulations in the plasma membrane. High levels of polyunsaturated PLs on the inner and outer sides of the plasma membrane would be beneficial for sustaining large membrane undulations, which are a series of convex and concave deformations. As a prerequisite for hemifusion, the decrease in cell thickness in the DHA-fed cells likely increases the probability of close apposition between the two undulating apical and basal membranes, explaining the increase in the frequency and the number of TEMs. Further experimental work on model membrane systems and coarse-grained simulations of TEM formation will help test this idea.

## Materials and Methods

### Reagents

LifeAct-GFP-pCMV plasmid was purchased from Ibidi. Antibodies used in this study were mouse anti-Na^+^/K^+^ ATPase (Santa Cruz), Annexin II (BD Transduction Laboratories), Calregulin (Santa Cruz), ERGIC 54 (Santa Cruz), and LAMP1 (BD Transduction Laboratories), and sheep anti-TGN46 (BioRad). Secondary Alexa Fluor-conjugated antibodies were from ThermoFisher and secondary peroxidase-conjugated antibodies were from Jackson ImmunoResearch. For immunofluorescence, hoechst 33342 and Alexa-fluor conjugated FITC-phalloidin were purchased from ThermoFisher. For STED imaging, Star635-phalloidin was purchased from Abberior. C3 toxin was purified as described (Boyer et al., 2006).

OA and DHA fatty acids (Sigma-Aldrich) were conjugated with fatty acid-free BSA (Sigma-Aldrich). Fatty acids were dissolved in warm (60°C) 200 mM NaOH and conjugated with BSA at the molar ratio of 5:1. The FA-BSA was aliquoted and filled with argon to minimalized oxidation. Lipoprotein depleted serum (LPDS) were prepared as described (Renaud et al., 1982). In brief, fetal bovine serum was loaded with NaBr to increase density to 1.21 g/ml followed by ultracentrifugation at 220,000g at 10°C for 48 hours in a Beckman Ti70 rotor. After centrifugation, a greasy layer containing lipoproteins appeared on the top of the tube was removed and the supernatant was centrifuged again at 220,000g at 10°C for 24 hours to remove the remaining lipoprotein. Later the serum was dialyzed intensively with Earle buffer (115 mM NaCl, 5.4 mM KCl, 1.8 mM CaCl_2_, 0.8 mM MgSO_4_, 5 mM Hepes, pH 7.4) in 14 kD cut-off dialysis membrane (Spectrum) for 72 hours and the buffer was refreshed for at least 5 times.

### Cell Culture, Treatment, and Transfection

HUVECs were cultured and electroporated, as described in (Stefani et al., 2017). In brief, HUVECs were trypsinized and suspended in Ingenio Solution (Mirus) containing the targeted DNA (10 g per 10^6^ cells) in a 4-mm cuvette (CellProjects). Then, cells were electroporated at 300 V, 450 µF, one pulse by GenePulser electroporator (BioRad).

To enrich HUVECs with OA or DHA, cells were washed twice with PBS and lipid starved in LPDS medium (Human endothelial SFM, 20% LPDS, 20 ng/ml FGF, 10 ng/ml EGF, 1 µg/ml Heparin, and Zellshield) overnight with or without 50 µg/ml ExoC3 toxin perpared as described in (Boyer et al., 2006) as indicated. Before experiments, cells were supplemented with 125 µM FA-BSA for 6 hr.

### Lipid extraction and Lipidomic

A clear description of lipidomic analysis was in supplementary material. Briefly, a modified Bligh and Dyer (Bligh and Dyer, 1959) extraction was carried out on cell pellets and purified cell membranes in order to extract lipids, which were then separated by chromatography with a C18 column and an appropriate gradient of mobile phase. The mass spectrometry analyzes were done using a Q-Exactive mass spectrometer (ThermoFisher) operating in data dependent MS/MS mode (dd-MS2) and the data were then processed using LipidSearch software v4.1.16 (ThermoFisher) in product search mode.

### Video Microscope

HUVECs were electroporated with LifeAct-GFP-pCMV as described above and seeded on gelatin coated polymer coverslip dish (Ibidi). After recovering for 24 hours from transfection, cells were lipid starved as in LPDS containing medium overnight. OA-BSA and DHA-BSA were added to the cells to the final concentration at 125 µM for 6 hours prior to video recording. Cells were supplemented with 25 mM Hepes (pH 7.4) and recorded on a 37°C heated stage of Nikon Ti inverted microscope using Ultraview spinning disk confocal system (Perkin Elmer). For the TEM opening, images were taken every 10 seconds for 1 hour. For TEM closure, images were taken every minute for 3 hour to avoid phototoxicity and bleaching during the acquisition. Acquired videos were analyzed by an Icy based automatic protocol. Acquired videos were analyzed by an Icy based automatic protocol.

### Image Analysis

Time-lapse videos were analysed with the Icy software (de Chaumont et al., 2012) and segmentation plugins (icy.bioimageanalysis.org/plugin). Each TEM was first manually identified as a region of interest (ROI). Considering the gradual recruitment of LifeAct-GFP around TEMs, it was difficult to properly identify the edge of TEMs. Indeed, the non-homogeneous contrast at the TEM border leads to a difficult clipping process. To overcome this challenge, we used advanced image analysis methods like the Active Contour plugin to properly track TEM over time. This allowed us in particular to determine the surface of the TEMs at each time point. We then applied a post-processing analysis to filter the TEMs and automatically eliminate remaining wrong segmentations. For instance we discarded any TEMs that display excessive growing air. In the end this protocol allowed us to provide precise statistics of the TEM dynamics, such as the evolution of the area, the diameters or the sphericity of TEMs over time.

### GPMV purification

Cells were enriched with OA or DHA as described above followed by induction of GPMV as described (Sezgin et al., 2012). Cells were washed with GPMV buffer (10 mM HEPES, 150 mM NaCl, 2 mM CaCl_2_, pH 7.4) twice and incubated with GPMV buffer containing 25 mM PFA and 2 mM DTT for 1 hour. Blebs were formed and released as GPMVs. Supernatant was collected and centrifuged at 100 g for 5 minutes to remove cell debris. Supernatant containing GPMVs was centrifuged in Beckman Type 70 Ti rotor at 20,000 g at 4 °C for 1 hr. GPMVs appeared as a transparent pellet and was collected for lipidomic analysis or western blot. For western blot, GPMV was lysed in Leammli buffer (50 mM Tris pH 7.4, EDTA 5mM, 2% SDS) and protein concentration was determined with BCA assay kit (Thermo Fisher) using BSA suspended in Leammli buffer as standard. GPMV lysate was adjusted to the same protein quantity. Glycerol, β-mercaptoethanol, and bromophenol blue were added to final concentration of 10%, 5% and 0.004%. Protein samples were analyzed by SDS-PAGE western blot.

### STED

Cells were grown on H1.5 glass coverslips coated with 10 µg/ml fibronectin. After treatment, cells were fixed with 4% PFA/0.1% glutaraldehyde for 15 minutes at room temperature. Cells were washed with PBS, quenched in 50 mM NH_4_Cl for 15 minutes followed by permeabilizing in IF buffer (PBS/0.05% saponin/0.2 % BSA) for 30 minutes. Later, the cells were stained with 1 µM Star635-phalloidin (Abberior) for 1 hr followed by 3 washes with IF buffer for 5 minutes and a final wash in H_2_O. The cells were mounted in Mount Solid Antifade (Abberior) following manufacturer’s instruction. Stimulated Emission Depletion (STED) imaging was performed by TCS STED SP8 (Leica) using a APO 93X/1.3 motCORR lens. The excitation laser was at 633 nm and pulse depletion laser at 775 nm. STED images were deconvolved using Huygens with 5 iterations.

### Atomic force microscope measurement of TEM topology

AFM experiments were carried out on a JPK NanoWizardIII mounted on a Zeiss Axio Observer.Z1. For elasticity measurements PFQNM-LC-A-Cal cantilevers (Bruker) were used with the SNAP calibration method (Schillers et al., 2017). The AFM was operated in QI mode on the cytoplasmic region of living cells to record a 10 µm^2^, 20×20 pixels map of force curves with 2 µm ramp length, 200 pN force trigger and 50µm/s tip velocity. Force maps were computed using in-house software (pyAF) for the fitting of the indentation up to 40 nm using Hertz model. For height measurements of TEMs Olympus AC40 cantilevers were used with the SADER calibration method (Sader et al., 2016). A QI map of 150×150 pixels bigger than the size of the TEM was recorded on 4 % PFA fixed cells with an 800-nm ramp and 100 µm/s tip velocity. We then computed the zero-force topography by determining the point of contact, and drew several profiles across the TEM to measure its diameter and the height of the cell at the border. This was all performed on the JPK analysis software.

### Physical model of TEM opening

In the physical model for TEM opening dynamics (Gonzalez-Rodriguez et al., 2012), the driving force for opening is given by

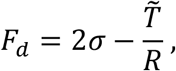

where σ is the membrane tension, 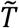 is the line tension and *R* is the TEM radius. The membrane tension σ depends on *R* through Helfrich’s law, which here we write in a generalized form to account for the coexistence of *N* simultaneous TEMs in the same cell:

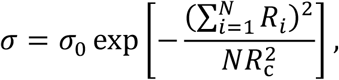

where 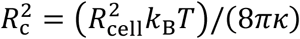 is the so-called critical radius, with *R*_cell_ the total cell radius, *k*_*B*_ the Boltzmann constant, *T* the temperature, and κ the membrane bending rigidity. Due to actin cable polymerization around the TEM, the line tension is not a constant but rather it increases with time, which can be represented by a linear increase 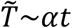 (Stefani et al., 2017). The dynamics of TEM opening are governed by a balance between driving force and cell-substrate friction, characterized by a friction coefficient μ. For the case of *N* identical TEMs, this balance results in the following differential equation:

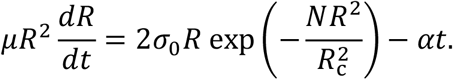

This equation can be solved numerically. However, insight can be gained by analytical approximations. First, in the limit of short time, when *R* is small, the equation can be approximated as

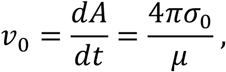

where *A*=π*R*^2^ is the TEM area. Therefore, TEM opening speed at short time, *v*_0_, is proportional to the undisturbed cell membrane tension σ_0_.

Second, the dependence of the maximum TEM area *A*_max_= π*R*max^2^ on the membrane parameters σ_0_ and κ can be estimated by the following approximation. Let us suppose that the initial opening speed *v*_0_ is an acceptable estimate of the average opening speed. Then, the opening time *t*_max_ is related to the maximum TEM area by

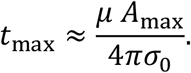

Moreover, at *t* = *t*_max_, the opening stops and d*R*/d*t*=0. By replacing these two results in the differential equation, we obtain the following approximate relationship:

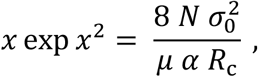

where we have defined *x* = *N R*_*max*_/*R*_*c*_. The nondimensional parameter on the right-hand side of this expression is slightly larger than 1, which requires *x* to be somewhat larger than 1. In this range of values, small changes in *x* yield large changes of the exponential function, implying that *x* is weakly dependent on the right-hand side. Therefore, *x* will remain approximately constant for moderate changes of σ_0_, implying that *NR*_max_ ∼ *R*_c_ ∼ 1/κ^1/2^. This result shows that the maximum TEM size is very sensitive to κ but rather insensitive to σ_0_. We thus obtain the following estimate of the membrane bending rigidity:

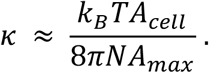

### Statistical analysis

Data are showed as the medium ± s.e.m. unless otherwise indicated. Data were analyzed with unpaired and two-tailed Mann-Whitney test unless otherwise indicated. P-value for *P<0.05, **P<0.01, ***P<0.001 and ****P<0.0001 were considered statistically significant. The statistical software used was Prism 8 (GraphPad Software, San Diego, CA, USA).

## Acknowlegdments

This work was supported by a grant from the Fondation pour la Recherche Médicale (Convention DEQ20180339156 Equipes FRM 2018), the Agence Nationale de la Recherche (ANR-11-LABX-0028-01, ANR-15-CE18-0016, ANR 10-EQPX-04-01, ANR- 10- INBS- 04, and ANR- 10- LABX-62-IBEID) and FEDER 12,001,407. MCT was supported by a PhD fellowship from the Labex Signalife PhD Programme and by the Fondation pour la Recherche Médicale (Contrat FDT201904008135). We thank Blandine Madji Hounoum for pilot lipidomics experiments. Frédéric Brau and Sophie Abelanet for support in light and STED microscopy. James Muncey for its helps in time-lapse analyses with Icy software.

## SUPPLEMENTARY FILES

### Supplementary Materials and Methods

A modified Bligh and Dyer (Bligh and Dyer 1959) was used for lipid extraction. One million cells were collected and pelleted in an eppendorf and 200 µL of water was added. After vortexing (30s), the sample was transferred in a glass tube containing 500 µL of methanol and 250 µL of chloroform. The mixture was vortexed for 30s and centrifuged (2500 rpm, 4°C, 10 minutes). Then, 300 µL of the organic phase was collected in a new glass tube and dried under a stream of nitrogen. The dried extract was resuspended in 60 µL of methanol/chloroform 1:1 (v/v) and transferred in an injection vial. The extraction protocol for purified cell pellet is the same as the one aforementioned in which every volume is divided by two.

Reverse phase liquid chromatography was selected for separation with an UPLC system (Ultimate 3000, ThermoFisher). Lipid extracts were separated on an Accucore C18 (150×2.1, 2.5µm) column (ThermoFisher) operated at 400 µl/ minutes flow rate. The injection volume was 3 µL. Eluent solutions were ACN/H_2_O 50/50 (V/V) containing 10mM ammonium formate and 0.1% formic acid (solvent A) and IPA/ACN/H_2_O 88/10/2 (V/V) containing 2mM ammonium formate and 0.02% formic acid (solvent B). The step gradient of elution was in %B: 0.0 min, 35%; 0.0-4.0 min, 35 to 60%; 4.0-8.0 min, 60 to 70%; 8.0-16.0 min, 70 to 85%; 16.0-25.0 min, 85 to 97%; 25-25.1 min 97 to 100% B, 25.1-31 min 100% B and finally the column was reconditioned at 35% B for 4 min. The UPLC system was coupled with a Q-exactive Mass Spectrometer (thermofisher, CA); equipped with a heated electrospray ionization (HESI) probe. This spectrometer was controlled by Xcalibur software (version 4.1.31.9.) and operated in electrospray positive mode.

Data were acquired with dd-MS2 mode at a resolution of 70 000 for MS and 35 000 for MS2 (at 200 m/z) and a normalized collision energy (NCE) of 25 and 30 eV. Data were reprocessed using Lipid Search 4.1.16 (ThermoFisher). The product search mode was used and the identification was based on the accurate mass of precursor ions and MS2 spectral pattern.

## Supplementary Figures

**SUP. FIGURE 1. Modification of neutral lipids during OA- or DHA-diet**.

(A) Fold change of neutral lipids compare to normal-cultured condition (FBS). TG triacylglycerol, DG diacylglycerol, Cer ceramide, SM sphingomyosine, ChE cholesterol ester. Data show means ± SEM; n>3 / experiments; 3 biological replicates. (B) The MS counts of each PC species normalized to total PC counts. (C) PI species distribution regarding their number of double bond (level of unsaturation): from zero to 6 double bonds as illustrated in figure 1B. AA and DHA contain 4 and 6 double bonds, respectively. LPDS: lipoprotein-depleted serum.

**SUP. FIGURE 2. Kinetics of OA and DHA-incorporation in HUVECs**.

(A) Kinetics of TG(18:1/18:1/18:1) formation in OA-treated cells (blue line). Kinetics of TG(22:6/22:6/22:6) formation in DHA-treated cells (pink line). (B) Kinetics of PUPL (orange line) and non PUPL (blue line) into the indicated lipid classes (PC, PE, PS, and PI) during OA treatment. (C) Kinetics of PUPL (orange line) and non PUPL (blue line) into PC, PE, PS, and PI during OA treatment. (B-C) PUPL contains at least one FA with at least 2 double bonds. Non-PUPL contains either SFA or MUFA on both acyl chains. (A-D) Data show means ± SEM; n>3 / experiments; 3 biological replicates.

**SUP. FIGURE 3. Impacts of C3 toxin on cellular lipidome**. (A) PL species distribution regarding their number of double bond (level of unsaturation): from zero to 6 double bonds as illustrated in figure 1B. AA and DHA contain respectively 4 and 6 double bonds. (B) Kinetics of DHA (red or pink line) and OA (blue or light blue line) incorporation into different lipid classes (PC, PE, PS, and PI) in the absence or presence of C3 toxin. Data show means ± SEM; n>3 / experiments; 1 biological replicates.

**SUP. FIGURE 4. Resume opening shows instability of TEM. (A) Scheme of resume opening**. TEM breaks locally and undergoes a second phase of opening. (B) Time-lapse images of TEM resume opening dynamics in OA- or DHA-fed cells using LifAct-GFP as label. (C-D) The evolution of TEM size over the period of recording in OA- (C) or DHA- (D) treated cells. (E) Correlation of TEM radius before (R_0_) and after resume opening (R_eq_). (F) Percentage TEM that undergo resume opening. (G) Difference in TEM radius (ΔR) before and after resume opening. (H) Speed of resume opening. (G-H) Data show median ± max to min of >5 TEMs. Data were analyzed with Mann-Whitney test and are not significant, ns.

**VIDEO 1. TEM tunnels have labile openings**. Video of LifeAct-GFP expressing HUVEC cells intoxicated with C3 exotoxin. The video was taken with spinning disk confocal at 1 frame per 10 seconds over the duration of 50 minutes. Scale bar, 20 µm.

**VIDEO 2. TEM tunnels in OA-treated cells**. Video of C3 intoxicated HUVEC cells treated with OA for 6 hours. LifeAct-GFP was transgenically expressed in HUVEC and the video was taken with a spinning disk confocal at 1 frame per 10 seconds over the duration of 50 minutes. Scale bar, 20 µm.

**VIDEO 3. TEM tunnels in DHA-treated cells**. Video of C3 intoxicated HUVEC cells treated with DHA for 6 hours. LifeAct-GFP was transgenically expressed in HUVEC and the video was taken with a spinning disk confocal at 1 frame per 10 seconds over the duration of 50 minutes. Scale bar, 20 µm.

