## Supplementary figures and images for "Docosahexaenoic fatty acid-containing phospholipids affect plasma membrane susceptibility to disruption by bacterial toxin-induced macroapertures"

### Suplemental figures

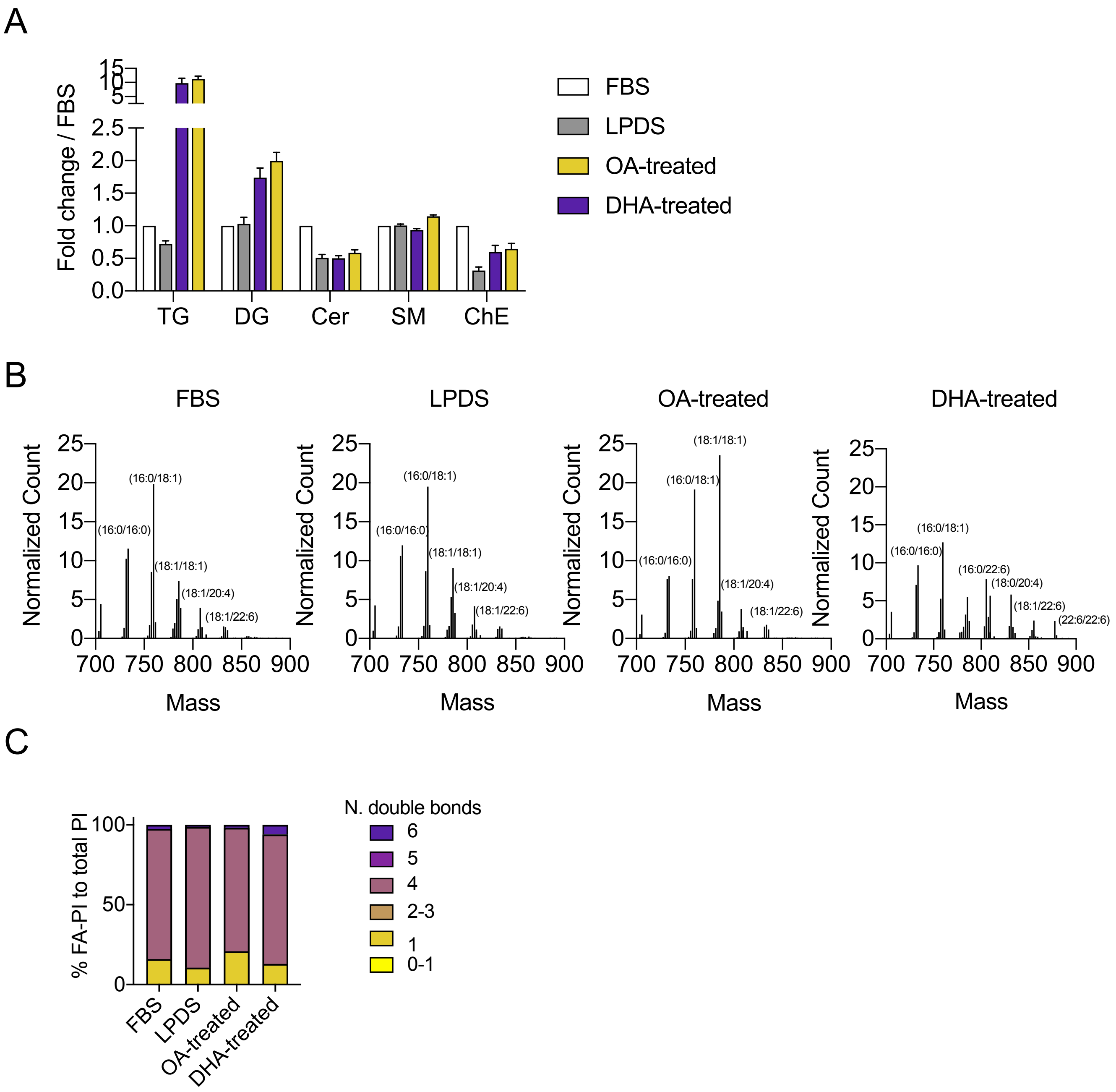

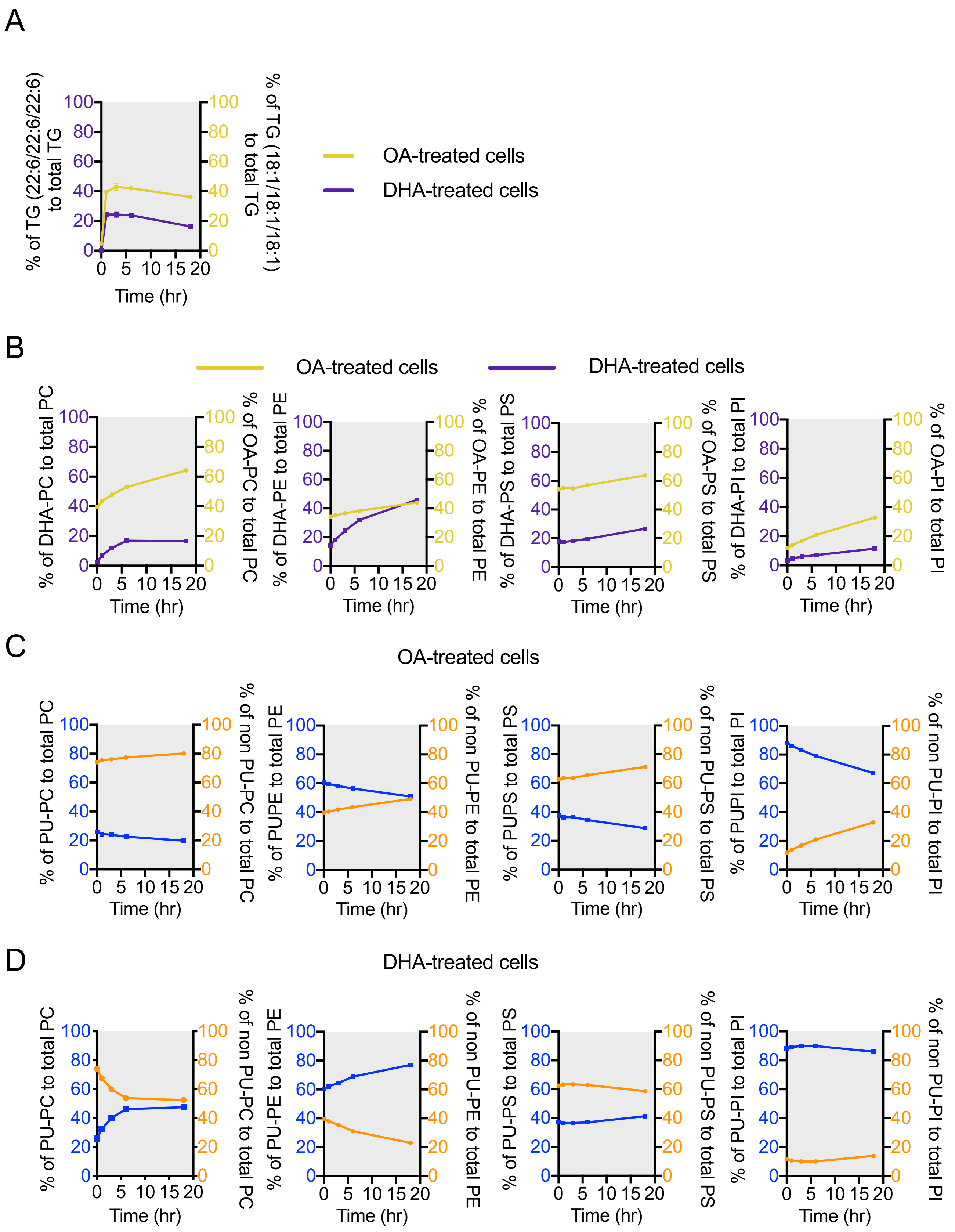

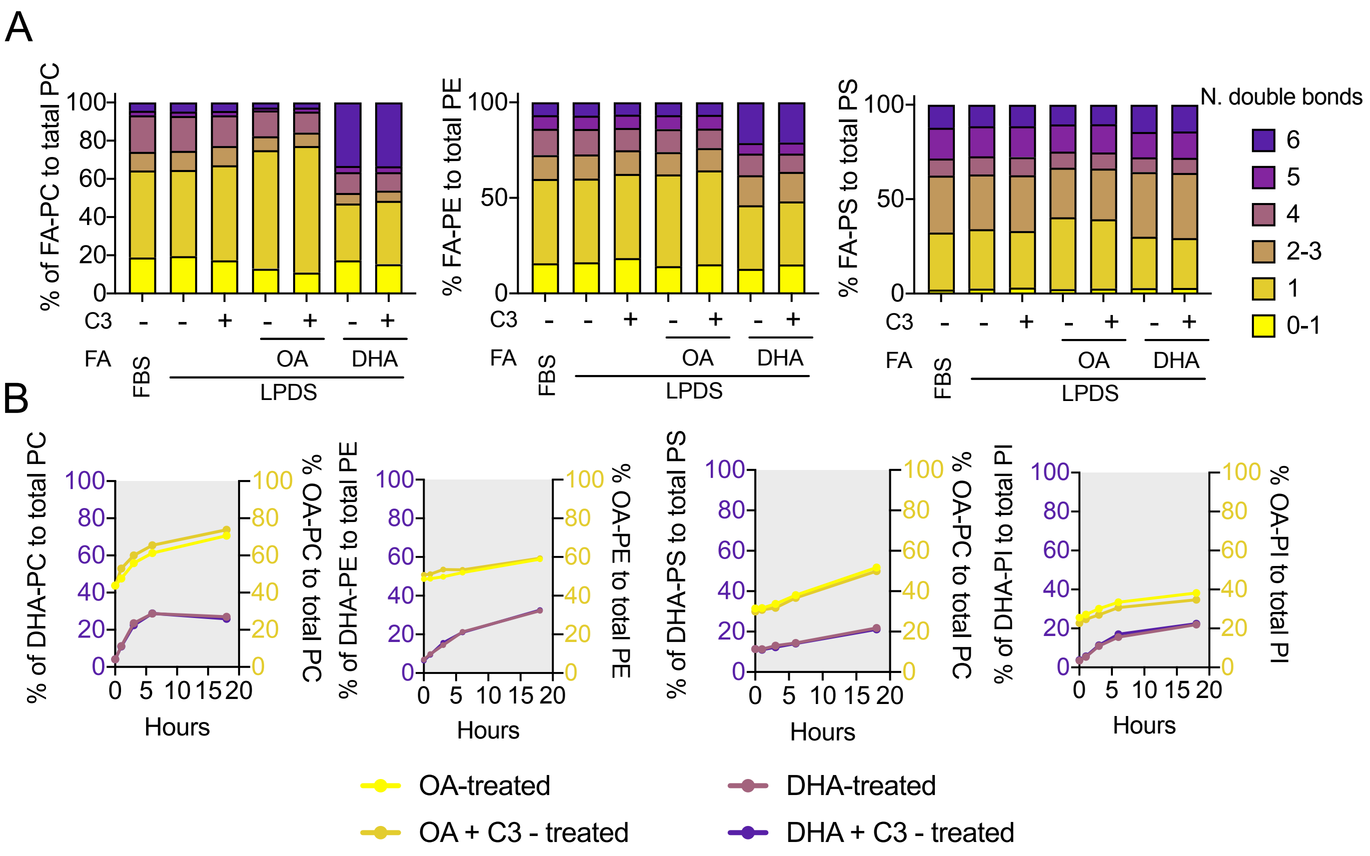

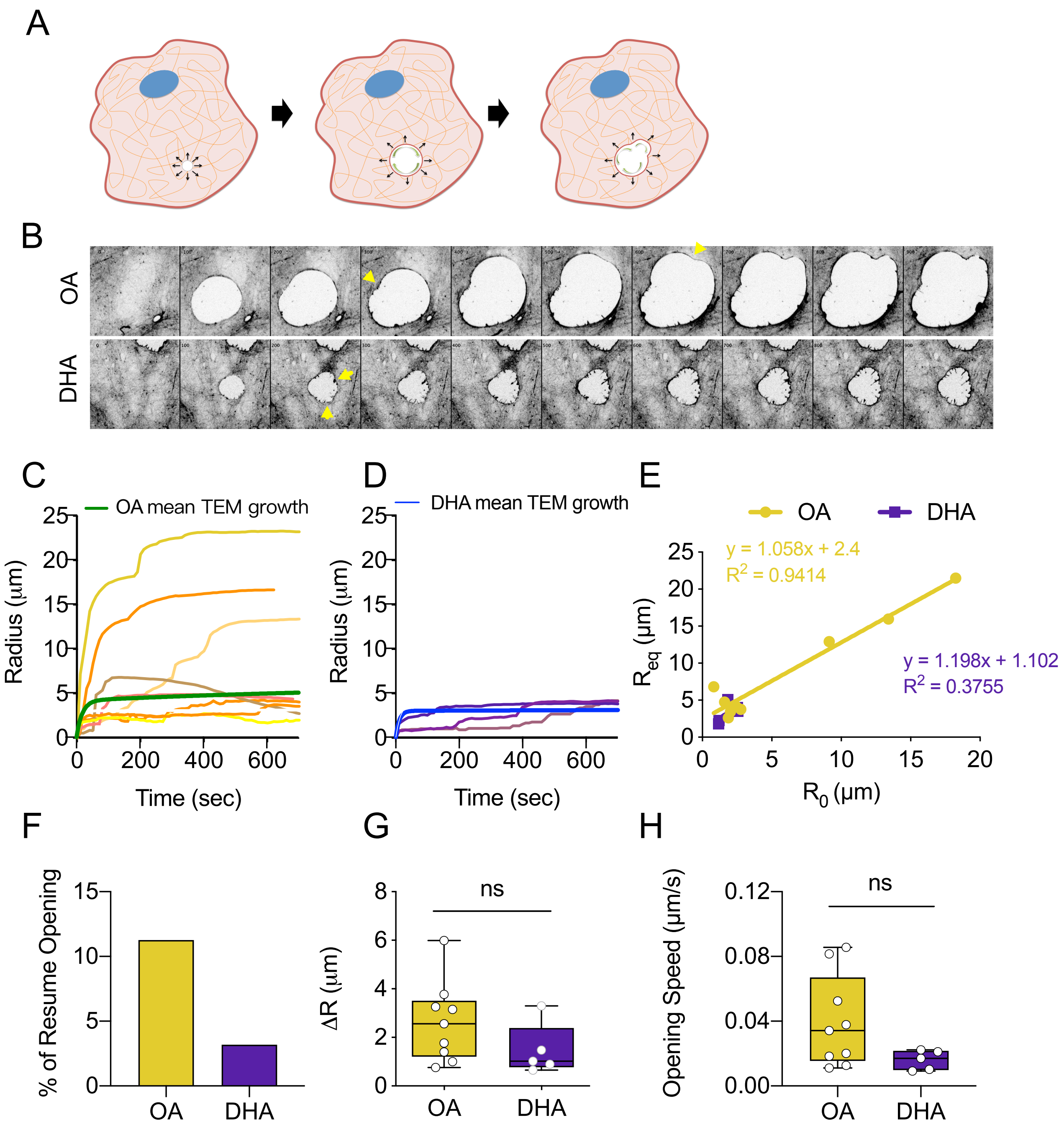
